# Predictive Dynamic Control Accurately Maps the Design Space for 2,3-Butanediol Production

**DOI:** 10.1101/2024.08.26.609681

**Authors:** Mathias Gotsmy, Anna Erian, Hans Marx, Stefan Pflügl, Jürgen Zanghellini

**Affiliations:** University of Vienna, Vienna, Austria; Austrian Centre of Industrial Biotechnology, Graz, Austria; BOKU University, Vienna, Austria; TU Wien, Vienna, Austria

**Keywords:** dynamic control, two-stage fed-batch process, two-reactor continuous process

## Abstract

2,3-Butanediol is a valuable raw material for many industries. Compared to its classical production from petroleum, novel fermentation-based manufacturing is an ecologically superior alternative. To be also economically feasible, the production bioprocesses need to be well optimized.

Here, we adapted and applied a novel process optimization algorithm, dynamic control flux-balance analysis (dcFBA), for 2,3-butanediol production in *E. coli*. First, we performed two-stage fed-batch process simulations with varying process lengths. There, we found that the solution space can be separated into a proportionality and a trade-off region.

With the information of the simulations we were able to design close-to-optimal production processes for maximizing titer and productivity, respectively. Experimental validations resulted in a titer of 43.6±9.9 g L^−1^ and a productivity of 1.93 ± 0.08 g L^−1^ h^−1^. Subsequently, we optimized a continuous two-reactor process setup for 2,3-butanediol productivity. We found that in this mode, it is possible to increase the productivity more than threefold with minor impact on the titer and yield.

Biotechnological process optimization is cumbersome, therefore, many processes are run in suboptimal conditions. We are confident that the methods presented here, will help to make many biotechnological productions economically feasible in the future.

**Highlights:**

- Precise simulations are used to sample the process solution space.
- Our simulations uncover big productivity potential in the 2,3-butanediol production.
- Experiments validate the predictions and show a 2,3-butanediol productivity improvement of 104 %.

## 1. Introduction

2,3-Butanediol is an important raw material in the chemical, pharmaceutical, cosmetics, agricultural, and food industries [2, 3, 4]. For example, it is used to produce synthetic rubber, fuel additives, perfumes, antifreeze agents, foods and pharmaceuticals [2, 4]. Fermentation-based 2,3-butanediol production is due to its economic and environmental sustainability an attractive alternative to petroleum-derived 2,3-butanediol [3].

There are several natural 2,3-butanediol producers, for example, from the genera *Klebsiella* [5, 6], *Enterobacter* [7, 8], and *Bacillus* [3]. They use 2,3-butanediol production pathway to prevent acidification [9], to regulate their internal NADH/NAD^+^ balance [10], and to store carbon [11].

Natural producers have several drawbacks. They often require complex growth media and may be pathogenic [1, 2]. Therefore, metabolic engineers have introduced genes for the 2,3-butanediol synthesis pathway into more commonly used organisms, such as *Lactobacillus lactis* and *Saccharomyces cerevisiae* [3]. For instance, Erian et al. [1] introduced three genes (*budaA, budB, budC*) from *Enterobacter cloacae* subsp. *dissolvens* into *E. coli* (indicated by red boxes in Figure 1).

**Figure 1.**
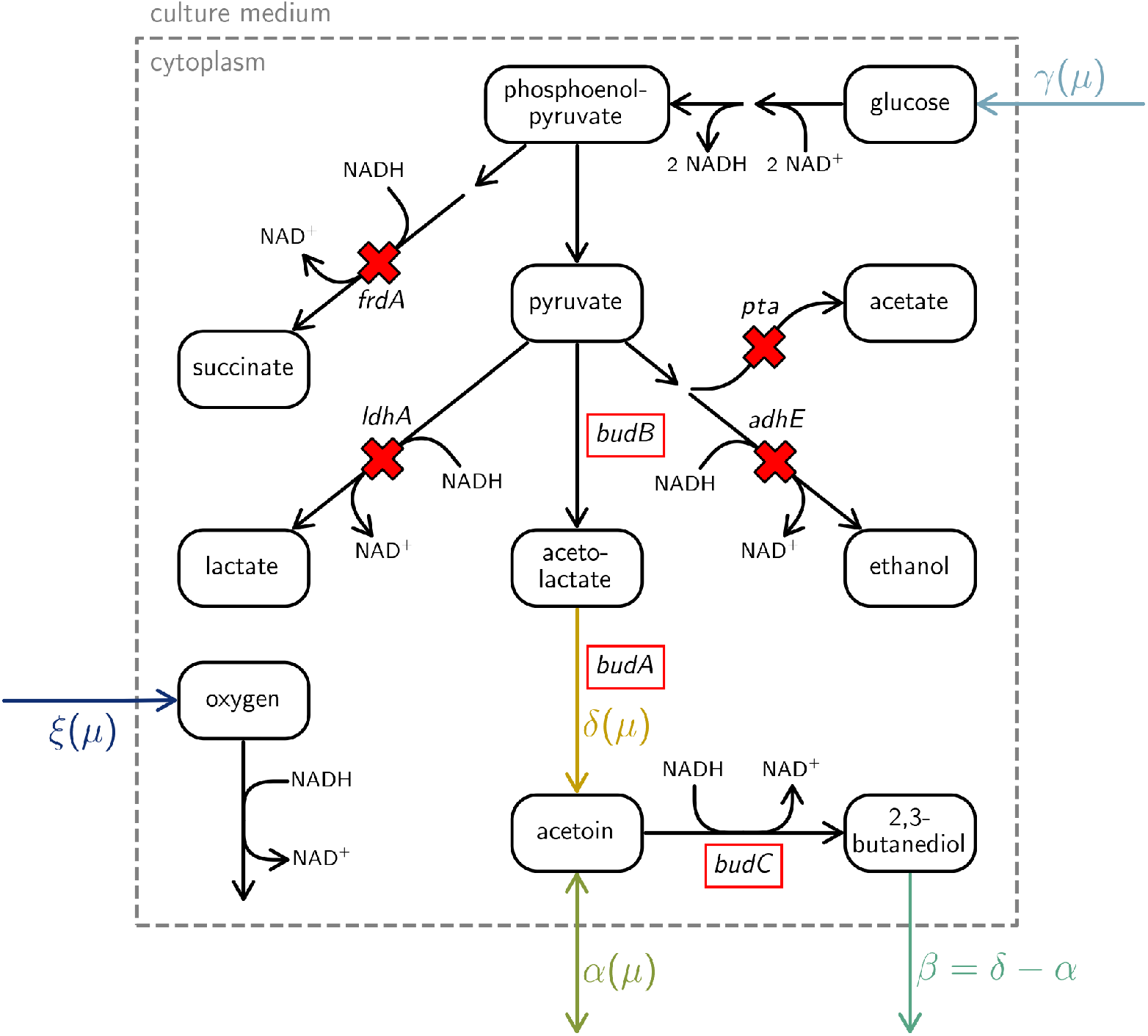
Selected metabolic reactions of the 2,3-butanediol production model of *E. coli* 445_EdissΔ4 [1]. Gene deletions (indicated by red crosses) and insertions (indicated by red boxes) were performed as described by Erian et al. [1]. In the resulting model, the NADH produced in the glycolysis can only be regenerated over respiration (requires a flux in *ξ*) or production of 2,3-butanediol (requires a flux in *β*). Broken arrows indicate multi-step pathways.

To make fermentation-based 2,3-butanediol a viable raw material for industry, large quantities must be produced quickly. Many studies focus on optimizing titer and yield, but productivity is equally important as it impacts the reactor size, which is a major investment cost in large-scale production plants [12]. As a rule of thumb, productivities below 2 g L^−1^ h^−1^ are considered uncommercializable [12].

Generally, 2,3-butanediol production rates are higher in microaerobic conditions, compared to fully aerobic ones [1]. This is due to the NAD^+^ regenerating properties of *budC* in the 2,3-butanediol production pathway (Figure 1). Conversely, growth rates are lower in microaerobic than in aerobic conditions. This may favor the design of two-stage processes for the best production conditions [13, 14, 15]. In these processes, a high cell density is achieved in an initial first stage, then bioprocess controls are switched to induce product formation in the second stage.

Additional methods for increasing the performance of 2,3-butanediol production processes include growth medium engineering [4], metabolic enzyme knockout and overexpression [1, 4], and enforced ATP wasting [16, 14]. Moreover, mathematical models of (genome scale) metabolic networks can help to uncover new strategies for increasing the production [17, 18]. For example, in a study for plasmid DNA production, theoretical analysis predicted a new method of growth decoupling via sulfate limitation [19]. Other methods comprise the discovery of optimal two-stage fermentations in batch processes [20], the optimization of feed rate and temperatures [21], the approximation of metabolic fluxes by neural networks in hybrid models [22], and network response analysis for rational strain engineering [23].

Another approach to improve the economic feasibility of biotechnological 2,3-butanediol production is the use of cheaper carbon sources. For example, optimization of sugar molasse medium compositions were performed for 2,3-butanediol producing strains of *Enterobacter ludwigii* [24] and *Vibrio natriegens* [25]. Moreover, downstream processing of the fermentation broth requires precise engineering [26].

In this study, we apply state-of-the-art process optimization algorithms to improve the production of 2,3-butanediol in *E. coli*. First, we construct a metabolic model of the high-2,3-butanediol producing strain 445_EdissΔ4 from Erian et al. [1]. Subsequent optimization elucidates the process solution space, which can be separated into a proportionality and a trade-off region. Furthermore, it predicts a big potential in increasing the productivity. Conducted validation experiments underline the precision of the simulations.

## 2. Methods

### 2.1. Theoretical Analysis

#### 2.1.1. Metabolic Model Construction

To perform our theoretical analysis, we first reconstructed a metabolic model that reflected the (engineered) pathways of the 2,3-butanediol-producing *E. coli* 445_EdissΔ4 from reference [1], using the *E. coli* core model from Orth et al. [27]. To adjust it to the genotype of the highly 2,3-butanediol-producing strain 445_EdissΔ4 [1], several reactions were deleted and introduced. To limit the possibility of anaerobic NAD^+^ regeneration, the genes *adhE, ldhA*, and *frdA* (corresponding to ethanol, lactate, and succinate production, respectively) were deleted. Additionally, *pta*, a gene of the acetate excretion pathway, is knocked out. Subsequently, reactions for the genes *budB, budA*, and *budC* representing the 2,3-butanediol production pathway were added to the metabolic model. Importantly, *budC* reintroduces the ability of anaerobic NAD^+^ regeneration via the reduction of acetoin to 2,3-butanediol [1]. A visualization of relevant metabolic reactions is given in Figure 1.

#### 2.1.2. Production Envelope Definition

We used experimental data from a previous study where *E. coli* 445_EdissΔ4 was grown in a two-stage (first aerobic, then microaerobic) process [1]. As these two conditions represent different metabolic states, following molecule uptake and secretion rates were fitted once per condition: glucose (*γ*), acetoin (*α*), 2,3-butanediol (*β*), and diols (*δ* = *α* + *β*) as well as the growth rate (*µ*). By setting the fitted rates as bounds of the metabolic model, the minimal oxygen uptake rate (|*ξ*|) for both conditions was calculated with flux balance analysis (FBA) [28] using CobraPy [29].

To create a continuous production envelope (PE) from the previously calculated feasible points we assumed a linear dependence of *γ*(*µ*) and *δ*(*µ*). The feasibility of all points along this line was verified with FBA.

#### 2.1.3. Fed-Batch Process Simulations

Process simulations were performed with an adapted algorithm for dynamic control flux balance analysis (dcFBA) [30]. dcFBA is a handy way of translating the bi-level optimization problem of dynamic flux balance analysis (dFBA) and process optimization into a single-level problem. This significantly improves convergence and results, especially in longer, more complex fed-batch simulations [31, 30].

##### Control Problem

Optimizing the dcFBA simulations involves estimating state variables from an initial state at time *t*_0_ = 0 throughout the process length (*T*). In this case study, the state variables are: biomass (*X*), 2,3-butanediol (*B*), acetoin (*A*), and the consumed glucose (*G*_con_). The differential equations of the state variables read

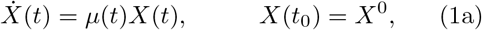

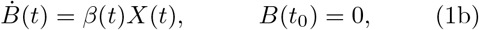

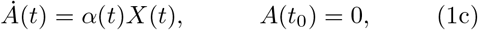

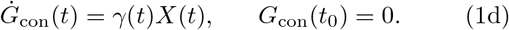

Here, 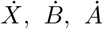, and *Ġ*_con_ represent the rates of change for biomass, 2,3-butanediol, acetoin, and consumed substrate, respectively. The parameters *µ, β* = *δ* − *α, α*, and *γ* denote the specific growth rate, production rates for 2,3-butanediol and acetoin, and substrate uptake rate.

The dynamic control problem derived from [30] reads: maximize the objective function *F* (*·*) over the control variable vector **µ** (i.e., a growth rate value per finite element of the simulation)

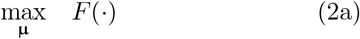

subject to (1) FBA constraints and FBA optimality constraints (i.e., KKT conditions constraints) per finite element, (2) moving finite elements length constraints, (3) orthogonal collocation of the differential equation, (4) a maximum of consumed glucose,

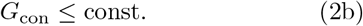

(5) a fixed process length,

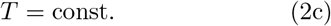

and (6) a upper and lower bound of growth rate control variables *µ* ∈ **µ**

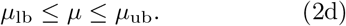

In the following paragraphs, parts of the control problem are explained in more detail. For even more information regarding the definition and optimization of dcFBA, we refer the readers to [30].

##### Process Target Metrics

In this study, we were interested in the optimization of two process performance metrics, the final product titer

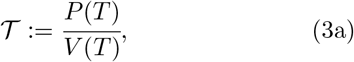

and the average productivity

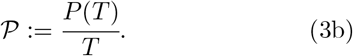

subject to the response of the cellular production host. Additionally, the product-to-substrate yield

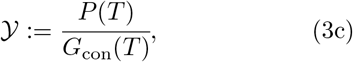

is often mentioned in literature [32].

##### Objective Function

Here, we adopted the objective function *F* of the dcFBA by adding a third term,

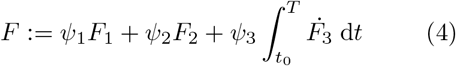

where *F*_1_ and *F*_3_ are one or a combination of state variables and *F*_2_ is the complementary slackness of the dual FBA formulation of the dcFBA [30]. Additionally, *ψ*_1_, *ψ*_2_, and *ψ*_3_ are scaling parameters. Concretely in this study, *F*_3_ = *P*, *ψ*_2_ = 10^1^, *ψ*_3_ = 10^−1^, and *ψ*_3_ ∈ {10^0^, 10^1^} for *F*_1_ ∈ {𝒯, 𝒫}, respectively.

##### Control Variables

We used the specific growth rate *µ* as control variable for the optimization. The integration of the differential equations was performed with orthogonal collocation [30] over *n*_FE_ = 30 finite elements. For each finite element, one value of *µ* was optimized. We emphasize that the control variable *µ* is a proxy for the oxygen uptake rate *ξ* (which is in practice controlled via the dissolved oxygen level). However, *µ, δ*, and *γ*, are linearly dependent on each other, while *ξ* is not (Figure 2). For the ease of optimization, we, therefore, chose to implement the PE in the dcFBA as a set of linear constraints. The value of *ξ* can be easily extracted from the PE and the corresponding optimized values of *µ*.

**Figure 2.**
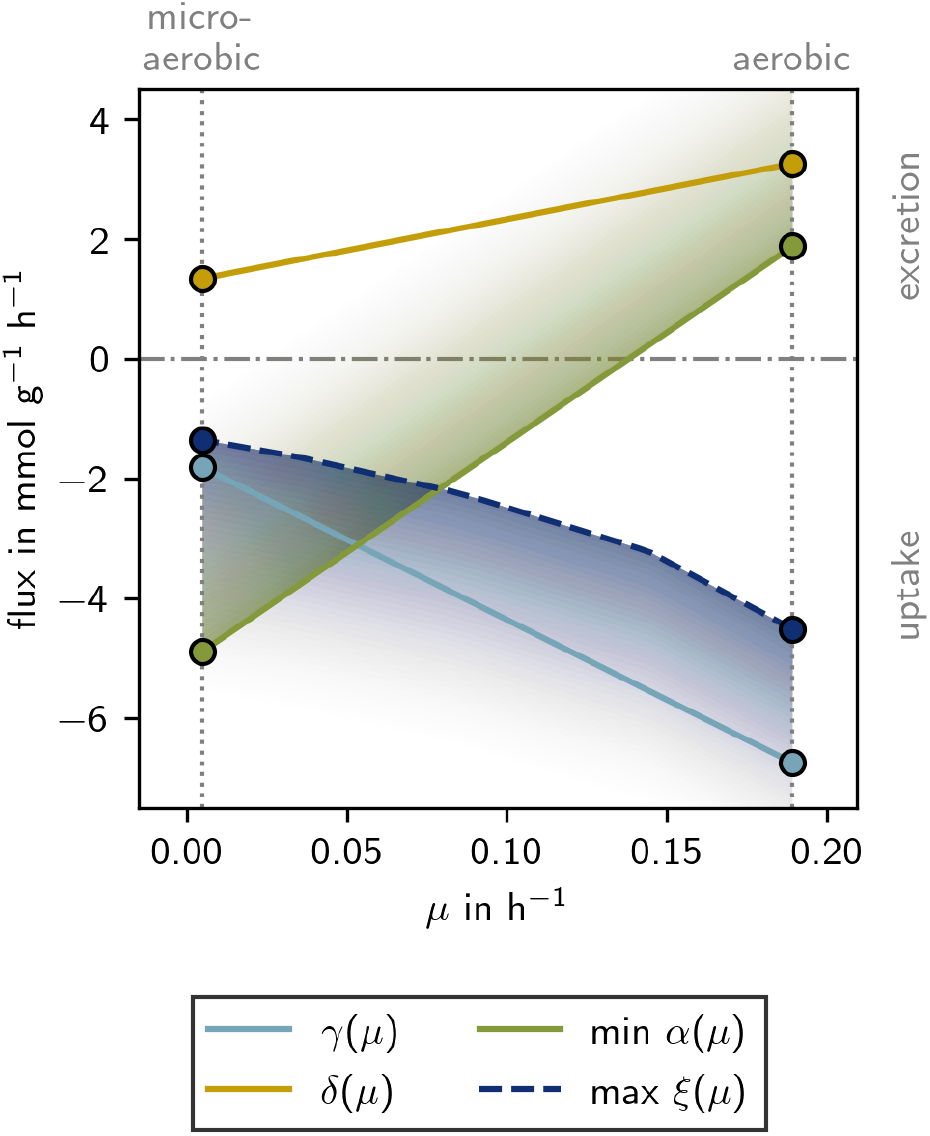
Fitted PE from the two-stage 2,3-butanediol production process reported in [1]. Markers indicate values fitted from the experimental data for the aerobic and microaerobic stage separately. Full lines represent weighted averages along *µ* between them, while the dotted line shows values calculated by FBA. As per convention, negative values indicate uptake from, and positive values indicate excretion into the culture medium.

##### Metabolic Model

Here, we use the stoichiometric matrix derived from the *E. coli* core model reconstructed in Section 2.1.1. Generally, dcFBA is fully compatible with using genome-scale metabolic models, however, we observed that the size of the stoichiometric matrix correlates with the required optimization time [30]. Therefore, if no core model is available, we recommend first identifying (exchange) rates of interest and then reducing the model size. This can be done by, for example, performing parsimonious FBA on different points of the production envelope and then discarding reactions that never carry any flux.

##### Process Solution Space

To map the process solution space, following constraints of the dcFBA were varied: *G*_con_ ∈ {200, 175} g, *µ*_lb_ between 0.005 and 0.189 h^−1^ in 11 equidistant steps, and *T* between 10 and 23.5 h in 10 equidistant steps and between 25 and 46 h in 8 equidistant steps. Moreover, we optimized for the titer 𝒯 and productivity 𝒫.

#### 2.1.4. Continuous Process Simulations

We optimized the steady state metabolic rates for two continuous reactors, as well as the feed flux (*ϕ*) and the glucose concentration in the feed (*G*_*ϕ*_).

##### Differential Equations

The differential equations for the continuous two-reactor setup read

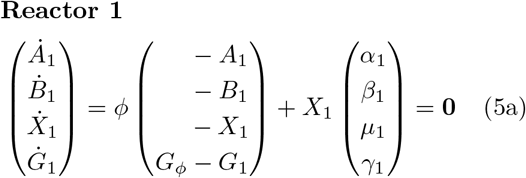

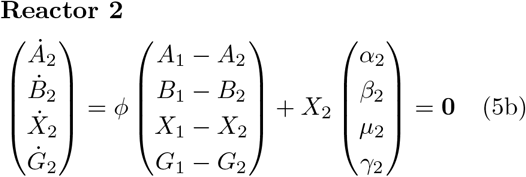

where the indices 1 and 2 represent the two reactors.

##### Objective Function

The objective function for the continuous two-reactor simulations read

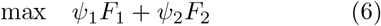

where *F*_1_ is one or a combination of state variables, *F*_2_ is the complementary slackness of the dual FBA formulation of the dcFBA and *ψ*_1_ and *ψ*_2_ are scaling parameters. Concretely in this study, *ψ*_1_ = 10^0^, *ψ*_2_ = 10^−2^, and *F*_1_ ∈ {*B*_2_, *ϕB*_2_} for titer and productivity optimization, respectively [33].

##### Numerical Integration

Subsequent to the optimization of the steady state, numerical integration of the continuous two-reactor setup was performed with SciPy [34]. The initial concentration of *X*_1_(*t* = 0) was set to the steady-state value, all other initial concentrations were set to 0.

### 2.2. Validation Experiments

To confirm the predictions of the dcFBA simulations, validation experiments were performed in duplicates.

#### 2.2.1. Upstream Process

Cultivations of *E. coli* W 445_EdissΔ4 were performed in duplicates analogous to Erian et al. (2018) [1] in a 1.2 L DASGIP Parallel Bioreactor system with 0.5 L working volume at 37 °C and pH7. After a batch with chemically defined medium containing 50 g L^−1^ glucose [1], cells were fed multiple times with a glucose-medium solution to approx. 50 g L^−1^ whenever glucose was depleted. Aerobic conditions in stage S1 were maintained by adjusting stirrer speed and aeration rates automatically to keep the dissolved oxygen level above 30 %. Microaerobic conditions (stage S2) were established by reducing the stirrer speed to 400 rpm and the aeration rate to 1 vvm. The switch from stage S1 to S2 was done for the control (CTL) and productivity optimized cultivations (MXP) at batch end (19.8 h, including lag phase) and with a 5 h delay (24.7 h, including lag phase), respectively.

#### 2.2.2. Analysis

Glucose, acetoin and 2,3-butanediol concentrations were determined by HPLC analysis (Shimadzu, Korneuburg, Austria) with an Aminex HPX-87H column (300 × 7.8 mm, Bio-Rad Laboratories, Hercules, CA) operated at 60 °C with 6 mM H_2_SO_4_ as mobile phase and a flow rate of 0.6 mL min^−1^ for 30 min. Peaks were detected and quantified with an RI detector or an UV lamp at 254 nm. To determine the biomass concentration, 2 mL culture broth was centrifuged for 5 min at 10 000 g at 4 °C, the pellet was washed once with deionized water and dried for 48 h at 100 °C. Biomass was determined in duplicates.

#### 2.2.3. Lag Phase Correction

For productivity calculation and comparison with simulations, we used a lag-free process length (*T*) which was estimated from the experimental process length (*T*_L_) as

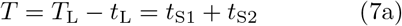

where *t*_S1_ and *t*_S2_ are the lengths of the (lag-free) aerobic stage I and the microaerobic stage 2. The lag phase length (*t*_L_) was calculated over an assumed exponential growth curve of the (aerobic) stage 1

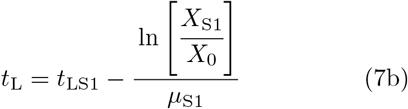

where *X*_S1_ and *X*_0_ is the biomass at the beginning and end of stage 1, *µ*_S1_ is the fitted exponential growth rate of stage 1, and *t*_LS1_ is the length of the uncorrected stage 1.

## 3. Results

We aim to design an optimal fermentation process for 2,3-butanediol production. Before doing so, we first need to characterize cellular behavior in the reference process (REF) [1]. In this process, 2,3-butanediol production occurs in two stages:

- Stage 1 (S1): Biomass is rapidly produced under aerobic conditions.
- Stage 2 (S2): Under microaerobic conditions, growth slows significantly, and the accumulated acetoin is rapidly converted into 2,3-butanediol. At the same time, de novo synthesis of 2,3-butanediol continues at a constant rate until the process is terminated.

### 3.1. Production Envelope

We analyze the experimental data for each state separately to determine uptake, secretion, and growth rates. In the constructed production envelope (PE) (Figure 2), we find that for each stage all rates remain constant over time (Supplementary Figure S1) with one exception: the acetoin exchange rate *α*. During the aerobic stage acetoin is constantly secreted (*α* > 0). However, when oxygen sparging throttles down, acetoin is initially consumed (*α* < 0) but depleted quickly (in ≤ 2.8 h) leading to a flux of *α* = 0 (Supplementary Figure S1K and L). Therefore, only a minimal value of *α* is shown in Figure 2.

We assumed that the transition between the aerobic and microaerobic flux states follows a straight line, characterized by linear relationships between *γ*(*µ*), min *α*(*µ*), and *δ*(*µ*) (the corresponding reactions are shown in Figure 1).

To check our model’s consistency, we used flux balance analysis (FBA) to calculate the minimally required oxygen uptake rate |*ξ*(*µ*)| (Figure 2). We find that oxygen uptake aligns with experimental observations [1]: the fitted rates from the microaerobic stage require less oxygen uptake than those from the aerobic stage. Since *ξ* is nonlinear, for further analyses, we use *µ* as the independent variable.

### 3.2. Fed-Batch Process Simulations

To further validate our fitted rates, we evaluated whether the PE defined above (Figure 2) could accurately replicate a real production process (i.e., REF) [1] *in silico*. For a fair comparison, we adjusted the experimental data to account for the observed lag time (*t*_L_), as detailed in the Methods Section (Supplementary Figure S2 and Table S1). This correction aligns the start of the simulation with the onset of exponential growth in the experimental data, ensuring precise comparison between the simulated and experimental results.

In Figure 3, we compare the lag-corrected experimental values (markers) with the process simulation (full line) of the reference process (REF) [1]. The close match between the simulations and experimental data gives us the confidence to proceed with further theoretical analyses.

**Figure 3.**
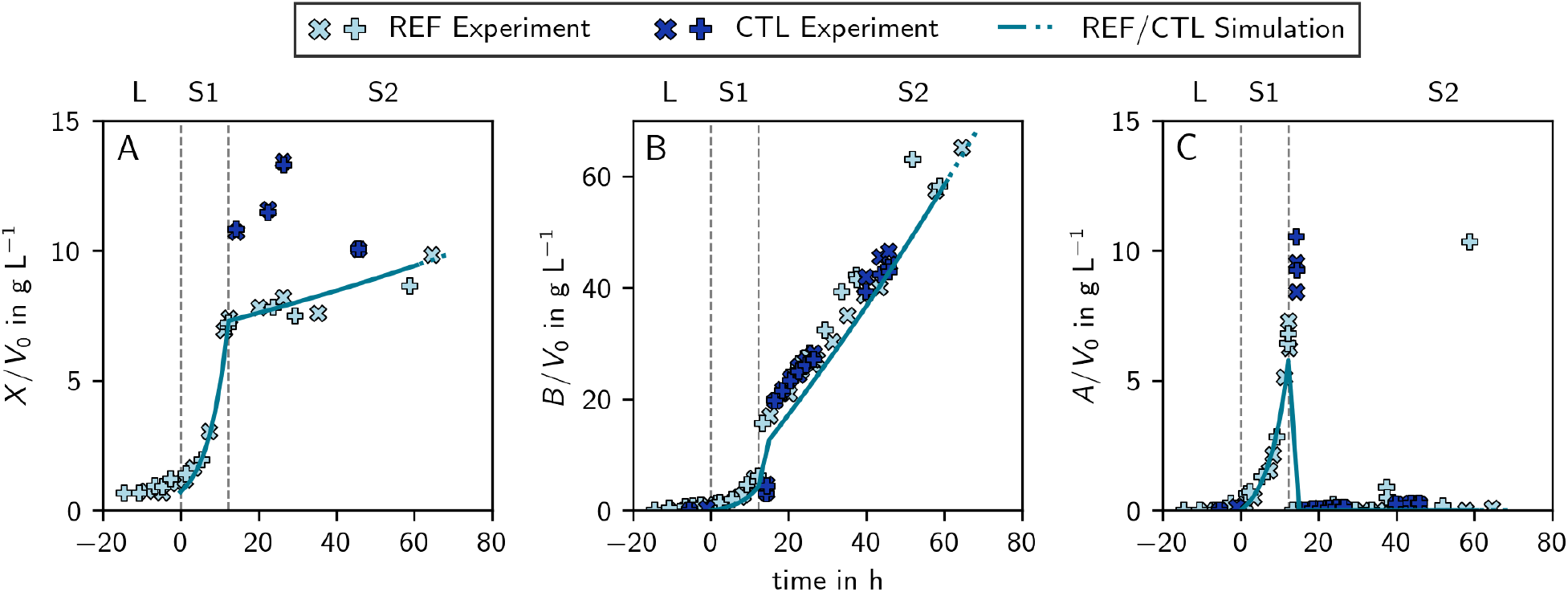
Simulation and experimental values of the reference (REF) and validation control (CTL) processes for biomass (A), 2,3-butanediol (B), and acetoin (C). The simulation is depicted as line (full and dotted) and experimental data as markers (different markers for duplicates). The difference between the REF and the CTL process is only the process length due to different max *G*_con_ (CTL: full line & *G*_con_ ≤ 175 g, REF: full and dotted line & *G*_con_ ≤ 200 g). The vertical dashed gray lines represent the borders of the three stages of the process: the lag phase (L), the aerobic stage (S1), and the microaerobic stage (S2). Experimental data was lag-corrected (Methods Section 2.2.3).

Next, we optimized the 2,3-butanediol production process using dynamic control flux balance analysis (dcFBA), with growth (serving as a proxy for oxygen consumption) as the control variable, subject to three major constraints:

(i) The total glucose consumption in each process was limited to at most 200 g, see Supplementary Figure S3.
(ii) The total process length (*T*) was fixed.
(iii) The minimal achievable specific growth rate (min *µ*) was fixed throughout the process.

Based on these settings, we computed and analyzed the maximum achievable productivity (𝒫) and titer (𝒯) as functions of process length and minimal growth rate (Figure 4A). The shaded (colors and gray) area characterizes feasible 2,3-butanediol production processes. The colored subspace indicates all processes where glucose is fully consumed. Colors indicate the minimal *µ* constraint, while dashed black lines represent the same *T* constraint. We mention that the zigzag inner boundary of the colored area is due to unavoidable sampling of the solution space and numerical instabilities at the edge of the feasible region.

**Figure 4.**
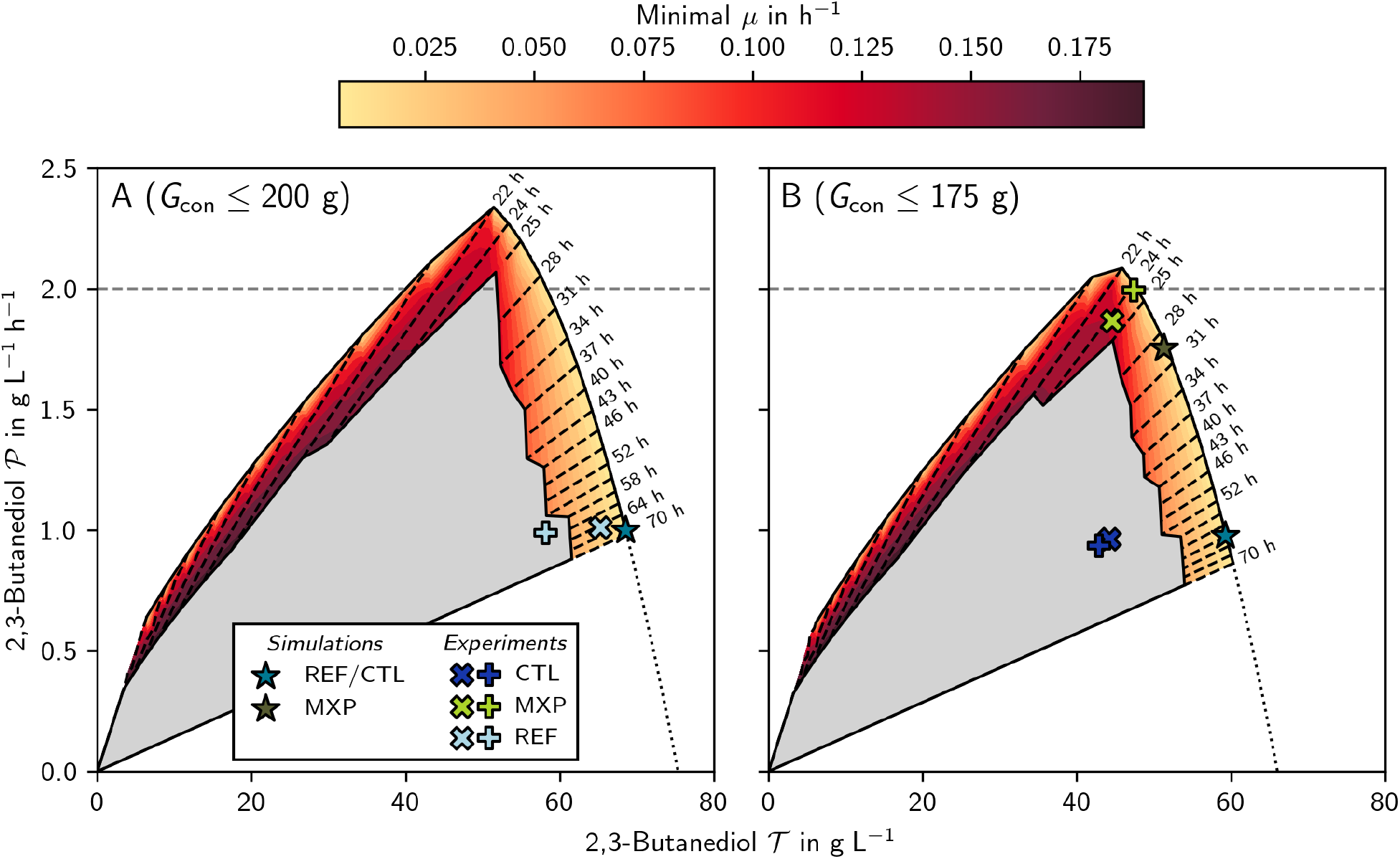
Optimal solution space of 2,3-butanediol production processes for *G*_con_ ≤ 200 g (panel A) and *G*_con_ ≤ 175 g (panel B). The shaded (colors and gray) area indicates feasible solutions and the colored subarea represents processes where all the glucose is consumed given a fixed process length (*T*, black dashed lines) and minimal growth rates (*µ*, colors). Plus and cross markers represent reference (REF, [1]) and validation (CTL, MXP) experiments. The star markers represent the results of the CTL, and MXP simulations. The dotted black line represents an extrapolation of the trade-off Pareto front (A: *y* = −0.00285*x*^2^ + 0.264*x* − 3.70, *R*^2^ *>* 0.99; B: *y* = −0.00319*x*^2^ + 0.254*x* − 2.84, *R*^2^ *>* 0.99).

In the following, we will focus our analysis on the colored area and make several observations:

(i) All optimal processes (except for poorly performing cases at the inside border of the solution space) are two-stage processes with constant growth rates in each stage. Biomass is rapidly produced under aerobic conditions, followed by a switch to microaerobic conditions for 2,3-butanediol production. Therefore, we can focus solely on the timing of the switch between them.
(ii) For a constant process length, both maximum productivity and titer increase as the minimal possible growth rate decreases.
(iii) For short processes (*T* ≤ 22 h), increasing the process duration enhances both maximum productivity and titer.
(iv) Maximum productivity is achieved for *T* = 22 h at the lowest possible minimal growth rate.
(v) For long processes (*T >* 22 h), maximum productivity decreases while the titer increases, resulting in a trade-off.
(vi) In the trade-off region (*T >* 22 h), productivity increases with the relative length of the first stage (*t*_S1_*/T*), assuming all glucose and acetoin are fully converted into 2,3-butanediol (Figure 5).
(vii) The maximum titer is reached with the largest simulated *T* = 70 h at the lowest possible minimal growth rate.
(viii) For fixed productivity, maximum titer depends non-monotonically on minimum growth rate (Supplementary Figure S4).
(ix) Variation the lower bound of *µ* (i.e., coloring of Figure 4) was done to investigate the sensitivity of the process with respect to *µ*. Interestingly, in the proportionality region sensitivity is small, however, in the trade-off region, it is considerable.

**Figure 5.**
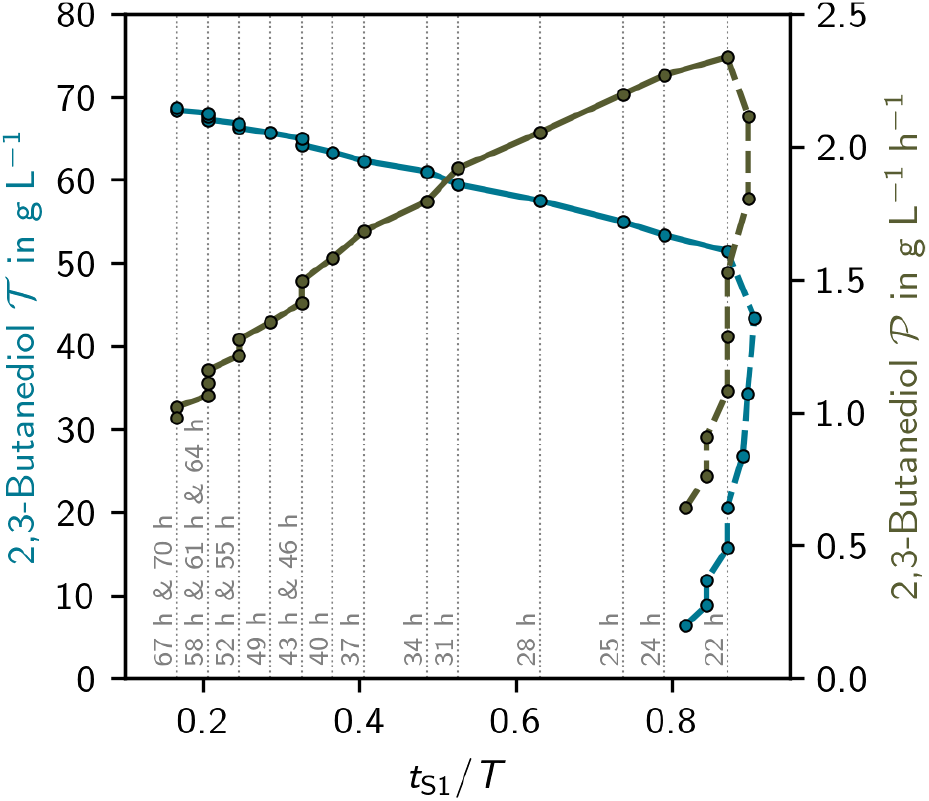
Influence of the relative first stage length (*t*_S1_*/T*) on the titer (*T*) and productivity (*P*) for min *µ* = 0.005 h^−1^. For processes in the trade-off region (*T* ≥ 22 h), *P* increases with the relative length of *t*_S1_*/T* and *T* decreases.

To further explore our observation (ii) “maximum productivity and titer increase as the minimal possible growth rate decreases”, we compare individual fermentations for a fixed process length of *T* = 22 h, as shown in Supplementary Figure S5.

When cells grow at the highest feasible minimal growth rate, they undergo essentially a one-stage growth process, resulting in the lowest productivity of 1.35 g L^−1^ h^−1^. In this scenario, biomass is rapidly produced from the available glucose, and 2,3-butanediol is generated in a growth-coupled manner without further conversion of the simultaneously produced acetoin.

For higher productivity, a two-stage process proves more effective because it utilizes the acetoin produced in the first stage. However, with a minimal *µ* of 0.134 h^−1^, acetoin is only partially converted to 2,3-butanediol (Supplementary Figure S5). Maximum productivity [observation (iv)] is achieved when all acetoin is fully converted to 2,3-butanediol, highlighting the advantage of a reduced growth rate that allows complete conversion.

To give more information about observation (vi) and Figure 5, we plotted Supplementary Figure S6. There, it is visible that although the total process length *T* increases with 2,3-butanediol titer 𝒯, the relative length of the first stage *t*_S1_*/T* is reduced.

### 3.3. Validation Experiments

In Figure 4A, the light blue markers represent the performance metrics of the reference process (REF) [1], which is also shown in Figure 3. This reference process operated near optimal conditions for high titer (Supplementary Figure S7). To validate our simulations, we explore the trade-off between productivity and titer and design and experimentally verify (in duplicates) a 2,3-butanediol production process that increases productivity rather than titer. As suggested by observation (vi), the length of the first stage (including the lag time, *t*_LS1_) was increased by 5 h resulting in the MXP process. The results of the validation experiments are shown in Figure 6 and 3 for the high productivity (MXP) and control (CTL) process, respectively. The process controls of MXP increased the average experimentally validated productivity by 104 % compared to CTL (Table 1).

**Table 1:** Overview of the process metrics of REF [1] and validation processes CTL and MXP. *X* was not measured at every time point, thus 𝒴 _*B/X*_ was calculated at adjacent time points given in the far right column.

| Run | Replicate | $T$<br>h | $\mathcal{T}$<br>$\text{g L}^{-1}$ | $\mathcal{P}$<br>$\text{g L}^{-1} \text{h}^{-1}$ | $\mathcal{Y}_{B/G_{\text{con}}}$<br>$\text{g g}^{-1}$ | $\mathcal{Y}_{B/G_{\text{con}}}$<br>$\text{mol mol}^{-1}$ | $\mathcal{Y}_{B/X}$<br>$\text{g g}^{-1}$ | $(T)$<br>(h) |
| --- | --- | --- | --- | --- | --- | --- | --- | --- |
| REF | A | 64.6 | 65.3 | 1.01 | 0.33 | 0.67 | 6.63 | ( 64.6) |
| REF | B | 58.8 | 58.2 | 0.99 | 0.29 | 0.57 | 6.73 | ( 58.8) |
| CTL | A | 45.9 | 44.2 | 0.96 | 0.24 | 0.49 | 4.62 | ( 45.7) |
| CTL | B | 45.9 | 42.9 | 0.93 | 0.25 | 0.50 | 4.33 | ( 45.7) |
| MXP | A | 23.8 | 44.5 | 1.87 | 0.26 | 0.52 | 2.45 | ( 23.8) |
| MXP | B | 23.8 | 47.4 | 1.99 | 0.27 | 0.54 | 2.70 | ( 23.7) |

**Figure 6.**
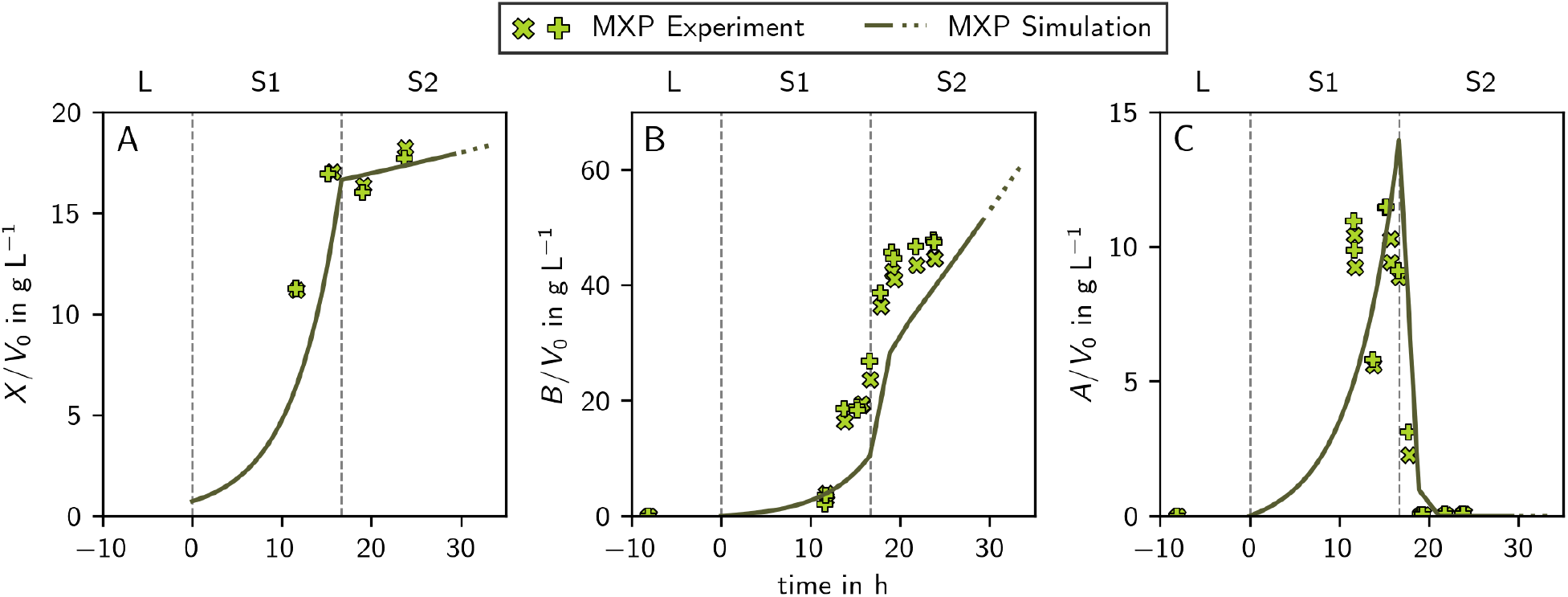
Simulation and experimental values of the MXP process for biomass (A), 2,3-butanediol (B), and acetoin (C). The simulation is depicted as line (full and dotted) and experimental data as markers (different markers for duplicates). The full line and markers indicate a process with *G*_con_ ≤ 175 g, the dotted line indicates an extrapolation to *G*_con_ ≤ 200 g. The vertical dashed gray lines represent the borders of the three stages of the process: the lag phase (L), the aerobic stage (S1), and the microaerobic stage (S2). Experimental data was lag-corrected (Methods Section 2.2.3).

In contrast to the optimal 𝒯 process, which was experimentally replicated very closely (REF, CTL), we designed our MXP more conservatively to the optimal 𝒫 predictions. The simulations showed a general improvement in productivity when extending *t*_S1_, however, in the optimum *t*_S2_ would be extremely short. As we were considering some biological delay which is not captured by the simulations, we settled on a slightly shorter *t*_S1_ (Supplementary Figure S7).

Due to bioprocess constraints, the validation experiments were performed with *G*_con_ ≤ 175 g. Therefore, we recalculated the process solution space for this amount of glucose, and compared it to 𝒯 and 𝒫 of CTL and MXP (Figure 4B).

Comparing REF (on which the simulations are based) and CTL validation experiments shows some deviation in the biomass concentrations (Figure 3A). This variation could be due to switching the process reactor system, which may have introduced an inconsistency.

Additionally, in Figure 6C, there is a drop in the acetoin concentration in stage 1 of the process. This drop coincides with a temporary exhaustion of glucose in the reactor before it is replenished again. We believe that during this brief period of glucose depletion, acetoin is consumed by the cells.

### 3.4. Continuous Process Simulations

To further increase the productivity of the 2,3-butanediol production process, we designed and simulated a continuous two-reactor bioprocess. In this setup, the aerobic and microaerobic stages occur simultaneously in two separate reactors, rather than being separated temporally. We optimized the process for productivity by using feed rate and glucose concentration (in the feed) as control variables while imposing constraints on the feasible biomass concentration and requiring complete glucose consumption. The result of this optimization is shown in Figure 7. The steady-state concentrations of all state variables and relevant process metrics are summarized in Supplementary Table S2 and Supplementary Table S3, respectively.

**Figure 7.**
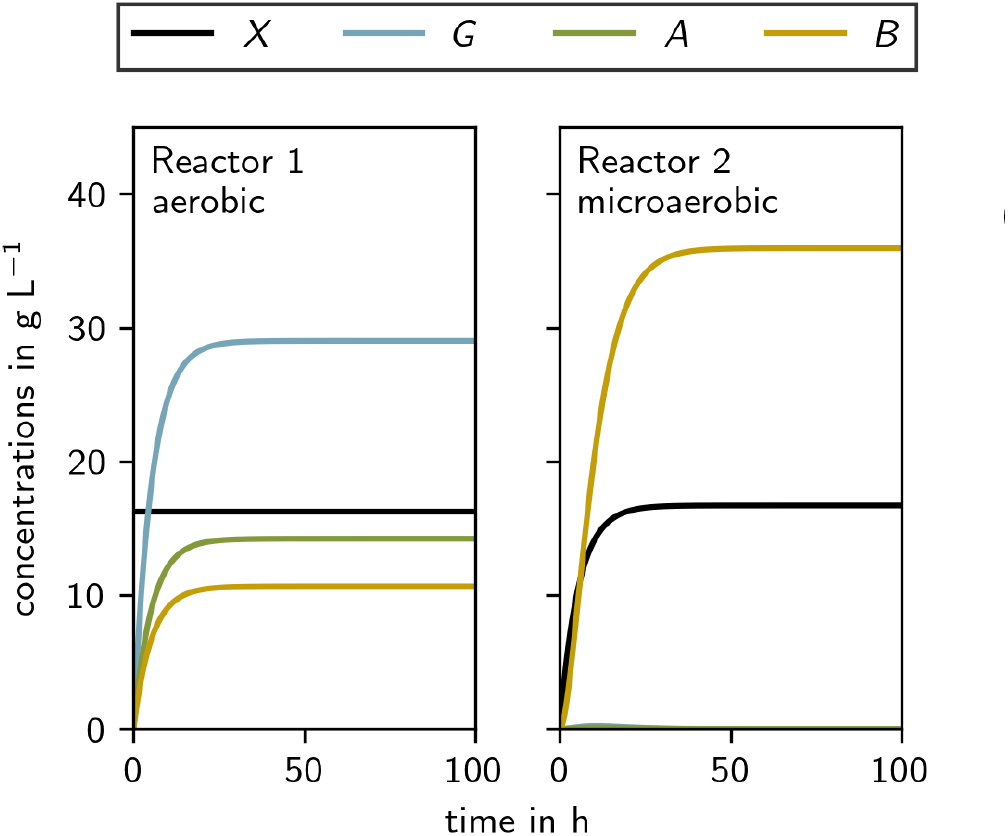
**Optimal productivity two-reactor continuous process simulation** where reactor A is run aerobically and reactor B is run microaerobically. The steady state concentrations are given in Supplementary Table S2 and the process metrics are given in Supplementary Table S3.

In the optimized two-reactor setup, the first reactor operates under aerobic condition, while the second operates under microaerobic condition. With a glucose feed rate of 25.2 g L^−1^ h^−1^, the simulated process could achieve microbial conversion to 2,3-butanediol with a titer of 35.9 g L^−1^ and a productivity of 6.80 g L^−1^ h^−1^. Even when constraining the titer to a minimal commercially viable value of 𝒯 ≥ 50 g L^−1^ as suggested by [12], our two-stage chemostat simulations predict more than double the productivity of a fed-batch process (𝒫 = 5.53 g L^−1^ h^−1^, Supplementary Figure S8 and Table S3.)

## 4. Discussion

Biobased 2,3-butanediol is a promising green platform chemical for various industries, but it faces significant economic hurdles compared to traditional petrochemical processes derived from crack gases. Although the petrochemical route is costly and energy-intensive [4], it remains competitive due to established infrastructure. In contrast, biobased 2,3-butanediol, though more environmentally friendly, struggles with high production costs driven by expensive fermentation media and complex processing. To be cost-competitive, biobased 2,3-butanediol production requires significant advancements in efficiency, scale, and cost reduction to replace petrochemical methods.

Building on our previous work [1], we developed and implemented a dynamic control algorithm [30] to optimize the feeding strategy and timing for a two-stage fed-batch process aimed at 2,3-butanediol production using a genetically modified *E. coli* strain. In the first stage, under aerobic conditions, cellular biomass increases rapidly while acetoin and 2,3-butanediol are simultaneously produced. During the transition to microaerobic conditions, the growth rate decreases, while 2,3-butanediol production increases to regenerate NAD^+^. In addition, the cells (re-)utilize any acetoin produced in the first stage and convert it into 2,3-butanediol.

Interestingly, although our dynamic control algorithm could continuously adjust the transition from aerobic to microaerobic conditions, the ideal process operates as a two-stage system: with maximum growth rate during the first stage and maximum acetoin uptake and 2,3-butanediol synthesis during the second stage. Within this design space, we find two specific optimal designs:

- The **titer-optimal** solution is characterized by a microaerobic stage that lasts as long as possible. In this scenario, the aerobic stage is designed only to ensure that all glucose is consumed. If all glucose can be consumed during the microaerobic stage, the process effectively becomes a single-stage process, with the titer- and yield-optimal solutions coinciding.
- The **productivity-optimal** solution is characterized by maximizing acetoin production during the aerobic phase, where biomass growth occurs at its highest rate and both acetoin and 2,3-butanediol are produced in a growth-coupled manner. In the subsequent microaerobic phase, 2,3-butanediol production is upregulated but further boosted by converting the accumulated acetoin into 2,3-butanediol. Maximum productivity is then achieved when all acetoin from the aerobic phase is fully converted to 2,3-butanediol in the microaerobic phase, aligned with the complete consumption of glucose. This approach maintains maximum 2,3-butanediol production for as long as acetoin is available, with productivity declining only after the acetoin is depleted.

Between these two optimal points, we observe a standard trade-off region, where increasing titer comes at the expense of reduced productivity, and vice versa [35]. This trade-off can be experimentally explored by varying the relative lengths of the two stages, a design parameter that our algorithm can accurately predict.

We remark that the productivity-optimal solution establishes a minimum required process length. For processes shorter than this minimum, both titer and productivity can be increased simultaneously; thus, such shorter processes are economically unappealing.

To further increase productivity, a promising strategy is to reduce the minimum required process length. This length is inversely related to the maximum growth rate and diol production rate, making the use of faster-growing cells essential. For example, employing fast-growing strains like *Vibrio natriegens* and *Bacillus licheniformis* has already yielded promising results [25, 36].

Chemostats are known for their high productivity. However, in our case, the growth and production phases occur under different conditions, requiring spatial separation in a multistage chemostat setup. Our simulations suggest that a two-stage chemostat could more than double productivity while maintaining high titers, potentially making the process truly economically viable [12]. However, these results may be overly optimistic, as challenges such as maintaining sterility during long runs and the risk of mutations were not considered, which continue to hinder the widespread adoption of continuous processes in biotechnological production [37].

Recently, we conducted a bioprocess optimization study using classical dynamic FBA (dFBA) for a two-stage fed-batch process [19]. Since dFBA was originally designed for process simulation rather than optimization, we had to employ a brute-force search to determine the optimal stage switching time points. In contrast, dcFBA enables the identification of the optimal switch time within given constraints in a single optimization run, significantly reducing computational time depending on the brute-force method’s resolution. Nevertheless, dcFBA was developed to result in a single optimized bioprocess. To uncover multi objective solution spaces (i.e., Fig. 4) additional algorithmic advances such as the normal boundary intersection method [38] may further reduce the computational cost required.

## 5. Conclusion

In this study, we apply a recently developed process optimization method, dynamic control flux balance analysis (dcFBA), to analyze the design space for optimal 2,3-butanediol production in fed-batch fermentation. Our simulations show that the optimal productivity-titer solution space consists of two distinct regions: a proportionality region and a trade-off region. In the proportionality region, both productivity and titer increase with longer process durations, but this region represents economically suboptimal processes. In contrast, the trade-off region is characterized by a decrease in productivity as process length increases, while titer continues to rise. At opposite ends of this region, we identify the titer-optimal and productivity-optimal processes, with the latter defining a minimum process length for economic viability. Our 2,3-butanediol production experiments conducted in duplicates validate these simulations.

Finally, simulations of continuous two-reactor fermentation indicate that further productivity gains are achievable.

Our study highlights the significant potential for improving bio-production processes and positions *in silico* process modeling as a powerful and reliable tool for efficiently exploring and optimizing process solution spaces, substantially reducing the effort and cost associated with traditional experimental methods.

## Author Contributions

Conceptualization: MG, JZ; Methodology: MG, AE; Software: MG; Validation: AE; Formal analysis: MG; Investigation: MG, AE; Resources: HM, SP; Writing - Original Draft: MG, JZ; Writing - Review & Editing: MG, JZ, AE, HM, SP; Visualization: MG; Supervision: JZ; Funding acquisition: JZ;

## Data Availability

All data and code for this study is available at https://github.com/Gotsmy/btdl.

## Footnotes

* *Preprint submitted to CSBJ*

## References

[1] A. M. Erian, M. Gibisch, S. Pflügl, Engineered E. coli w enables efficient 2, 3-butanediol production from glucose and sugar beet molasses using defined minimal medium as economic basis, Microbial Cell Factories 17 (2018) 1–17.

[2] J. W. Lee, Y.-G. Lee, Y.-S. Jin, C. V. Rao, Metabolic engineering of non-pathogenic microorganisms for 2, 3-butanediol production, Applied Microbiology and Biotechnology 105 (2021) 5751–5767.

[3] C. W. Song, J. M. Park, S. C. Chung, S. Y. Lee, H. Song, Microbial production of 2, 3-butanediol for industrial applications, Journal of Industrial Microbiology and Biotechnology 46 (2019) 1583–1601.

[4] S. Maina, A. A. Prabhu, N. Vivek, A. Vlysidis, A. Koutinas, V. Kumar, Prospects on bio-based 2, 3-butanediol and acetoin production: Recent progress and advances, Biotechnology advances 54 (2022) 107783.

[5] X.-J. Ji, H. Huang, J.-G. Zhu, L.-J. Ren, Z.-K. Nie, J. Du, S. Li, Engineering klebsiella oxytoca for efficient 2, 3-butanediol production through insertional inactivation of acetaldehyde dehydrogenase gene, Applied Microbiology and Biotechnology 85 (2010) 1751–1758.

[6] C. Ma, A. Wang, J. Qin, L. Li, X. Ai, T. Jiang, H. Tang, P. Xu, Enhanced 2, 3-butanediol production by klebsiella pneumoniae sdm, Applied microbiology and biotechnology 82 (2009) 49–57.

[7] M.-Y. Jung, H.-M. Jung, J. Lee, M.-K. Oh, Alleviation of carbon catabolite repression in enterobacter aerogenes for efficient utilization of sugarcane molasses for 2, 3-butanediol production, Biotechnology for Biofuels 8 (2015) 1–12.

[8] L. Li, K. Li, Y. Wang, C. Chen, Y. Xu, L. Zhang, B. Han, C. Gao, F. Tao, C. Ma, et al., Metabolic engineering of enterobacter cloacae for high-yield production of enantiopure (2r, 3r)-2, 3-butanediol from lignocellulose-derived sugars, Metabolic engineering 28 (2015) 19–27.

[9] J.-L. Tsau, A. A. Guffanti, T. J. Montville, Conversion of pyruvate to acetoin helps to maintain ph homeostasis in lactobacillus plantarum, Applied and environmental microbiology 58 (1992) 891–894.

[10] L. Johansen, K. Bryn, F. Stormer, Physiological and biochemical role of the butanediol pathway in aerobacter (enterobacter) aerogenes, Journal of Bacteriology 123 (1975) 1124–1130.

[11] Z. Xiao, P. Xu, Acetoin metabolism in bacteria, Critical reviews in microbiology 33 (2007) 127–140.

[12] S. Van Dien, From the first drop to the first truckload: commercialization of microbial processes for renewable chemicals, Current opinion in biotechnology 24 (2013) 1061–1068.

[13] S. Klamt, R. Mahadevan, O. Hädicke, When do two-stage processes outperform one-stage processes?, Biotechnology journal 13 (2018) 1700539.

[14] S. Boecker, B.-J. Harder, R. Kutscha, S. Pflügl, S. Klamt, Increasing atp turnover boosts productivity of 2, 3-butanediol synthesis in Escherichia coli, Microbial Cell Factories 20 (2021) 1–12.

[15] K. Shabestary, S. Klamt, H. Link, R. Mahadevan, R. Steuer, E. P. Hudson, Design of microbial catalysts for two-stage processes, Nature Reviews Bioengineering (2024) 1–17.

[16] F. Yatabe, T. Seike, N. Okahashi, J. Ishii, F. Matsuda, Improvement of ethanol and 2, 3-butanediol production in saccharomyces cerevisiae by atp wasting, Microbial Cell Factories 22 (2023) 204.

[17] F. Delvigne, R. Takors, R. Mudde, W. van Gulik, H. Noorman, Bioprocess scale-up/down as integrative enabling technology: from fluid mechanics to systems biology and beyond, Microbial biotechnology 10 (2017) 1267–1274.

[18] J. M. Park, H. Song, H. J. Lee, D. Seung, Genomescale reconstruction and in silico analysis of klebsiella oxytoca for 2, 3-butanediol production, Microbial Cell Factories 12 (2013) 1–11.

[19] M. Gotsmy, F. Strobl, F. Weiß, P. Gruber, B. Kraus, J. Mairhofer, J. Zanghellini, Sulfate limitation increases specific plasmid dna yield and productivity in E. coli fed-batch processes, Microbial Cell Factories 22 (2023) 242.

[20] K. Raj, N. Venayak, R. Mahadevan, Novel two-stage processes for optimal chemical production in microbes, Metabolic Engineering 62 (2020) 186–197.

[21] G. Schloegel, R. Lueck, S. Kittler, O. Spadiut, J. Kopp, J. Zanghellini, M. Gotsmy, Optimizing bioprocessing efficiency with optfed: Dynamic nonlinear modeling improves product-to-biomass yield, Computational and Structural Biotechnology Journal (2024).

[22] N. Adebar, S. Arnold, L. M. Herrera, V. N. Emenike, T. Wucherpfennig, J. Smiatek, Physics-informed neural networks for biopharmaceutical cultivation processes: Consideration of varying process parameter settings, Biotechnology and Bioengineering (2024).

[23] S. Tsouka, M. Ataman, T. Hameri, L. Miskovic, V. Hatzimanikatis, Constraint-based metabolic control analysis for rational strain engineering, Metabolic Engineering 66 (2021) 191–203.

[24] O. Psaki, S. Maina, A. Vlysidis, S. Papanikolaou, A. M. de Castro, D. M. Freire, E. Dheskali, I. Kookos, A. Koutinas, Optimisation of 2, 3-butanediol production by enterobacter ludwigii using sugarcane molasses, Biochemical Engineering Journal 152 (2019) 107370.

[25] A. M. Erian, P. Freitag, M. Gibisch, S. Pflügl, High rate 2, 3-butanediol production with vibrio natriegens, Bioresource Technology Reports 10 (2020) 100408.

[26] J. Haider, M. A. Qyyum, A. Hussain, M. Yasin, M. Lee, Techno-economic analysis of various process schemes for the production of fuel grade 2, 3-butanediol from fermentation broth, Biochemical Engineering Journal 140 (2018) 93–107.

[27] J. D. Orth, R. M. Fleming, B. Ø. Palsson, Reconstruction and use of microbial metabolic networks: the core Escherichia coli metabolic model as an educational guide, EcoSal plus 4 (2010) 10–1128.

[28] J. D. Orth, I. Thiele, B. O. Palsson, What is flux balance analysis?, Nat Biotech 28 (2010) 245–248. doi:10.1038/nbt.1614.

[29] A. Ebrahim, J. A. Lerman, B. O. Palsson, D. R. Hyduke, Cobrapy: constraints-based reconstruction and analysis for python, BMC systems biology 7 (2013) 1–27

[30] M. Gotsmy, D. Giannari, R. Mahadevan, J. Zanghellini, Optimizing fed-batch processes with dynamic control flux balance analysis, IFAC-PapersOnLine (2024).

[31] R. D. de Oliveira, G. A. Le Roux, R. Mahadevan, Non-linear programming reformulation of dynamic flux balance analysis models, Computers & Chemical Engineering 170 (2023) 108101.

[32] J. Nielsen, J. D. Keasling, Engineering cellular metabolism, Cell 164 (2016) 1185–1197.

[33] Y. L. Gordeeva, Y. A. Ivashkin, L. Gordeev, Modeling the continuous biotechnological process of lactic acid production, Theoretical Foundations of Chemical Engineering 46 (2012) 279–283.

[34] P. Virtanen, R. Gommers, T. E. Oliphant, M. Haberland, T. Reddy, D. Cournapeau, E. Burovski, P. Peterson, W. Weckesser, J. Bright, et al., Scipy 1.0: fundamental algorithms for scientific computing in python, Nature methods 17 (2020) 261–272.

[35] K. Zhuang, L. Yang, W. R. Cluett, R. Mahadevan, Dynamic strain scanning optimization: an efficient strain design strategy for balanced yield, titer, and productivity. dyssco strategy for strain design, BMC biotechnology 13 (2013) 1–15.

[36] L. Li, C. Chen, K. Li, Y. Wang, C. Gao, C. Ma, P. Xu, Efficient simultaneous saccharification and fermentation of inulin to 2, 3-butanediol by thermophilic bacillus licheniformis atcc 14580, Applied and environmental microbiology 80 (2014) 6458–6464.

[37] A. L. Zydney, Perspectives on integrated continuous bioprocessing—opportunities and challenges, Current Opinion in Chemical Engineering 10 (2015) 8–13.

[38] Y. I. Lim, P. Floquet, X. Joulia, Efficient implementation of the normal boundary intersection (nbi) method on multiobjective optimization problems, Industrial & engineering chemistry research 40 (2001) 648–655.

