## Supplementary Material for "Predictive Dynamic Control Accurately Maps the Design Space for 2,3-Butanediol Production"

*Supplementary Tables*

Table S1: Lag phase correction of 2,3-butanediol production processes from [1] and this study. All values of the (un-)corrected process length ($T_L$ and $T$) and aerobic stage 1 length ($t_{LS1}$ and $t_{S1}$) are given in hours. Surprisingly, there are big differences in the lag time length ($t_L$) for REF.

| | | uncorrected | | corrected | | |
|---|---|---|---|---|---|---|
| process | replicate | $t_{LS1}$ | $T_L$ | $t_L$ | $t_{S1}$ | $T$ |
| REF | A | 21.2 | 73.4 | 8.8 | 12.4 | 64.6 |
| REF | B | 26.8 | 73.4 | 14.5 | 12.2 | 58.8 |
| CTL | A | 19.8 | 51.3 | 5.5 | 14.3 | 45.9 |
| CTL | B | 19.8 | 51.3 | 5.4 | 14.4 | 45.9 |
| MXP | A | 24.7 | 31.9 | 8.0 | 16.7 | 23.8 |
| MXP | B | 24.7 | 31.9 | 8.1 | 16.6 | 23.8 |

Table S2: Steady state values of the state variables of the continuous process simulations in $\mathrm{g\,L^{-1}}$.

| | opt. $\mathcal{P}$ | | opt. $\mathcal{P}$, $\mathcal{T} \geq 50\,\mathrm{g\,L^{-1}}$ | |
|---|---|---|---|---|
| Variable | Reactor 1 | Reactor 2 | Reactor A | Reactor B |
| $X$ | 16.2 | 16.7 | 15.9 | 16.7 |
| $G$ | 29.0 | 0.0 | 49.6 | 0.0 |
| $A$ | 14.2 | 0.0 | 0.0 | 0.0 |
| $B$ | 10.7 | 35.9 | 31.6 | 50.0 |

Table S3: Process variables of the continuous process simulations.

| | $\phi$ $\mathrm{L\,L^{-1}\,h^{-1}}$ | $G_\phi$ $\mathrm{g\,L^{-1}}$ | $\mathcal{T}$ $\mathrm{g\,L^{-1}}$ | $\mathcal{P}$ $\mathrm{g\,L^{-1}\,h^{-1}}$ | $\mathcal{Y}_{B/G_{con}}$ $\mathrm{g\,g^{-1}}$ | $\mathcal{Y}_{B/G_{con}}$ $\mathrm{mol\,mol^{-1}}$ | $\mathcal{Y}_{B/X}$ $\mathrm{g\,g^{-1}}$ |
|---|---|---|---|---|---|---|---|
| opt. $\mathcal{P}$ | 0.19 | 133 | 35.9 | 6.80 | 0.27 | 0.54 | 2.15 |
| opt. $\mathcal{P}$, $\mathcal{T} \geq 50\,\mathrm{g\,L^{-1}}$ | 0.11 | 170 | 50.0 | 5.53 | 0.29 | 0.59 | 2.99 |

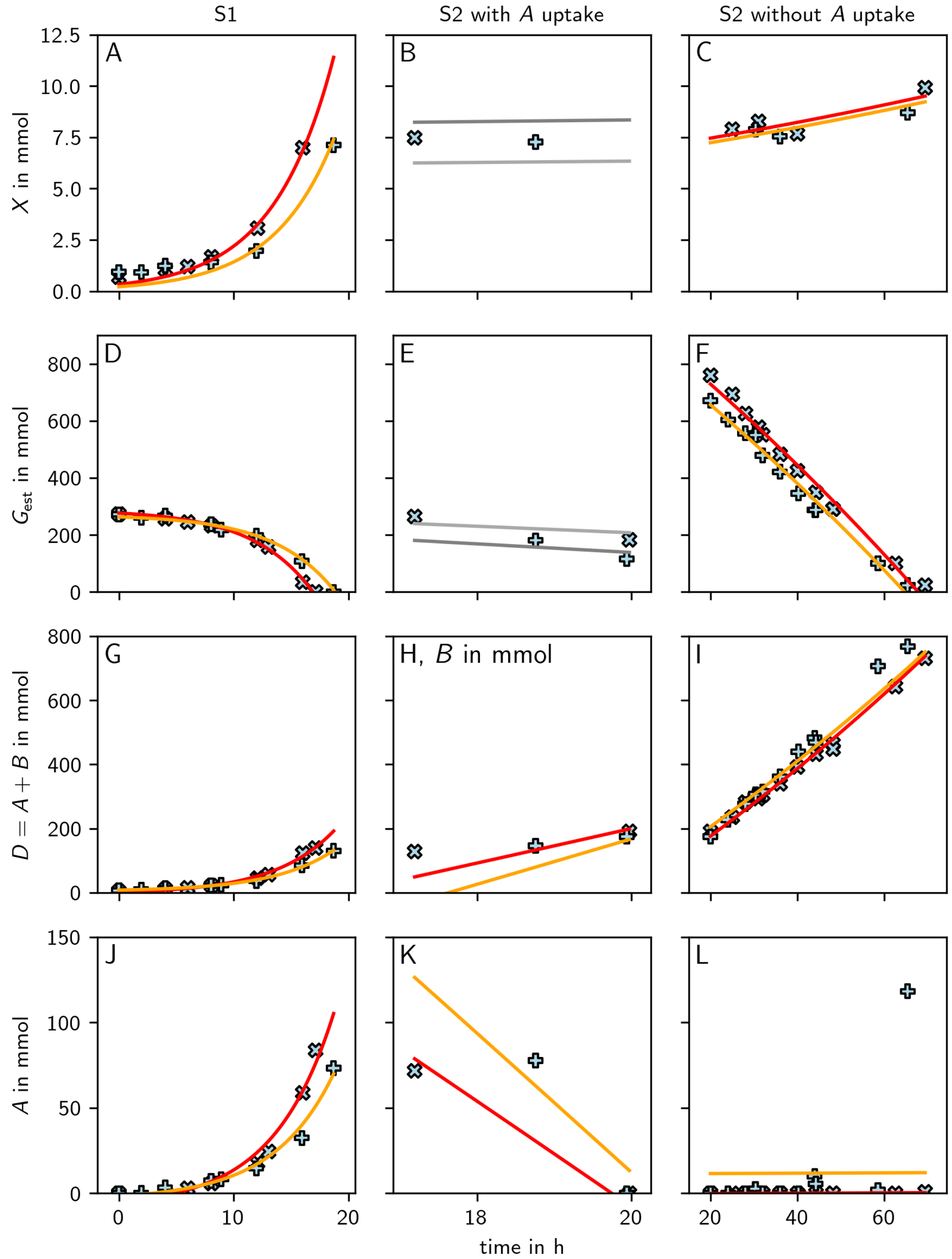


Figure S1: **Fitted rates of the aerobic (S1, first column) and microaerobic (S2, second and third column, with and without acetoin ($A$) uptake) stage.** The markers represent the experimental values of REF [1] and the lines show the fits. S1 and S2 were fitted independently. As acetoin uptake in the microaerobic stage was very short, initially the rates were fitted from time points where acetoin was depleted (third column). The estimated values of $\mu_{S2}$ and $\gamma_{S2}$, where then fixed (indicated as gray lines) for fitting of $\alpha_{S2}$ uptake (second column). $\beta_{S2}$ was fitted (Panel H), however, not actually used in the PE (Figure 2).

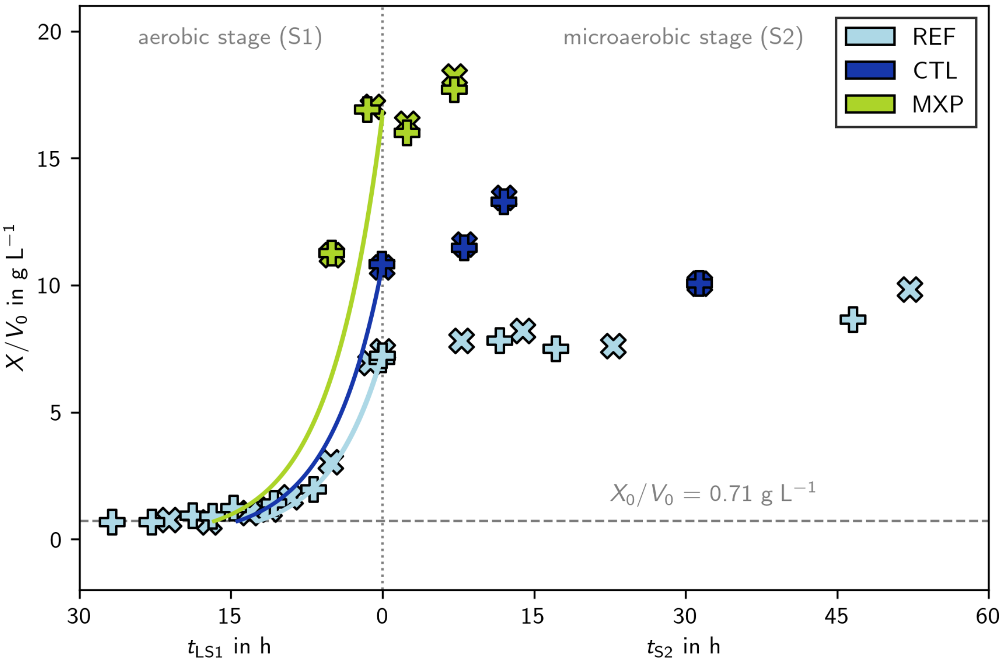


Figure S2: **A lag phase correction was performed prior to the productivity calculation.** The biomass concentration at the end of the aerobic first stage ($X_{\mathrm{LS1}}/V_0$) was estimated from a linear approximation of adjacent experimental values, and a initial biomass concentration ($X_0/V_0$) of $0.71\,\mathrm{g\,L^{-1}}$ was assumed. The corrected aerobic stage length ($t_{\mathrm{S1}}$) was calculated as the length of an exponential growth from the initial biomass concentration to the estimated biomass at $t_{\mathrm{S1}}$ with the $\mu_{t_{\mathrm{S1}}}$ estimated from the aerobic control processes.

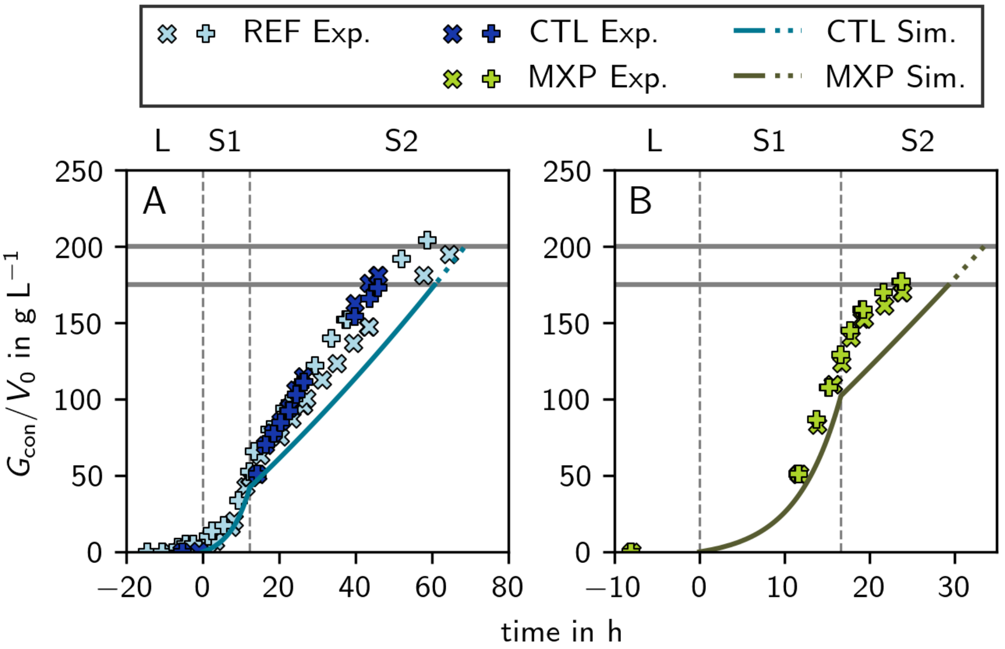


Figure S3: **Consumed glucose ($G_{\mathrm{con}}$) of the REF, CTL, and MXP experimental values and simulations.** The horizontal gray lines represent 200 g and 175 g glucose, respectively. The simulations until 175 g glucose are depicted as full lines and as dotted lines thereafter.

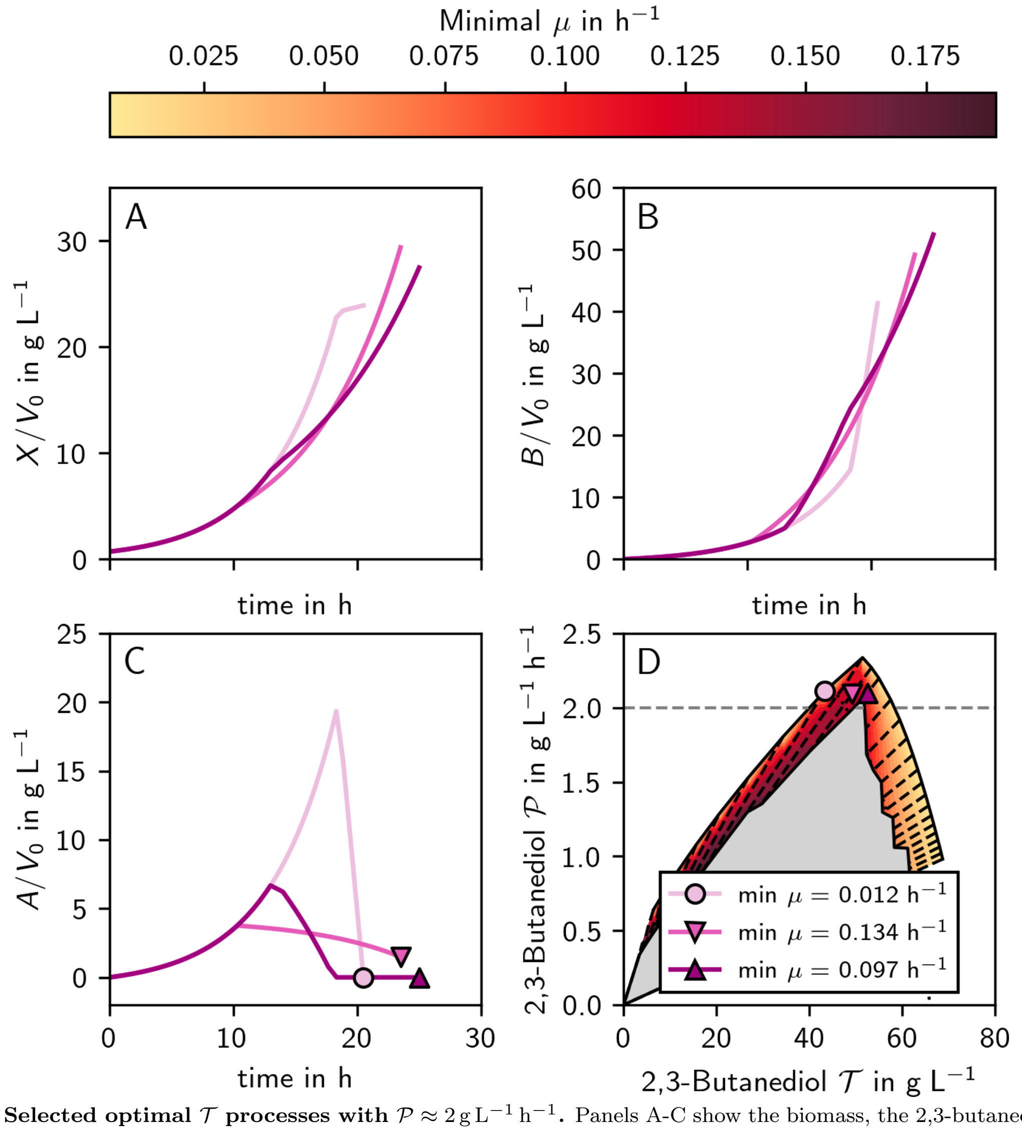


Figure S4: **Selected optimal $\mathcal{T}$ processes with $\mathcal{P} \approx 2\,\mathrm{g\,L^{-1}\,h^{-1}}$.** Panels A-C show the biomass, the 2,3-butanediol and the acetoin concentrations, panel D indicates the location of the selected process on the 2,3-butanediol production solution space.

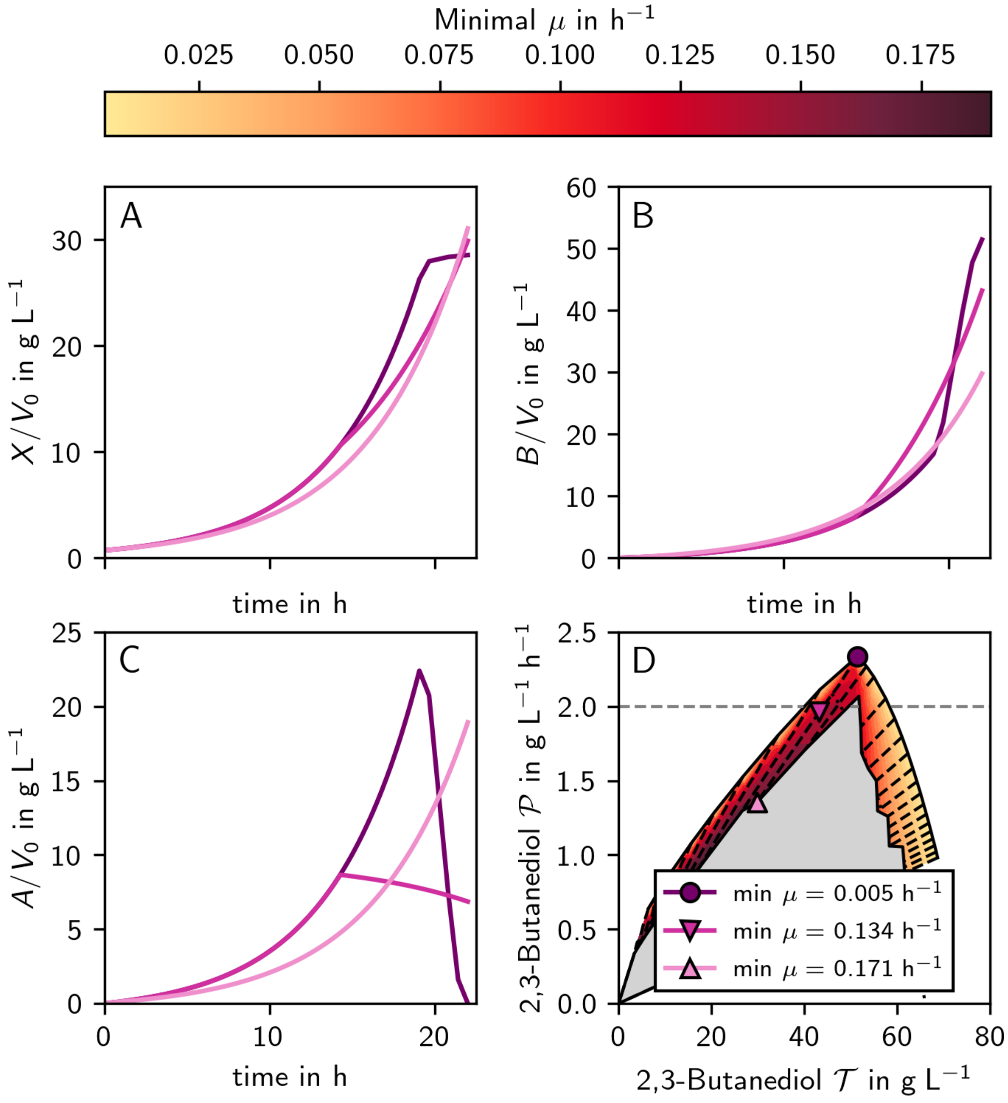


Figure S5: **Selected optimal $\mathcal{T}$ processes for $T = 22\,\text{h}$.** Panels A-C show the biomass, the 2,3-butanediol and the acetoin concentrations, panel D indicates the location of the selected process on the 2,3-butanediol production solution space. When $\min \mu$ becomes too large, not all $A$ can be converted into $B$ severely reducing the performance (Panel C).

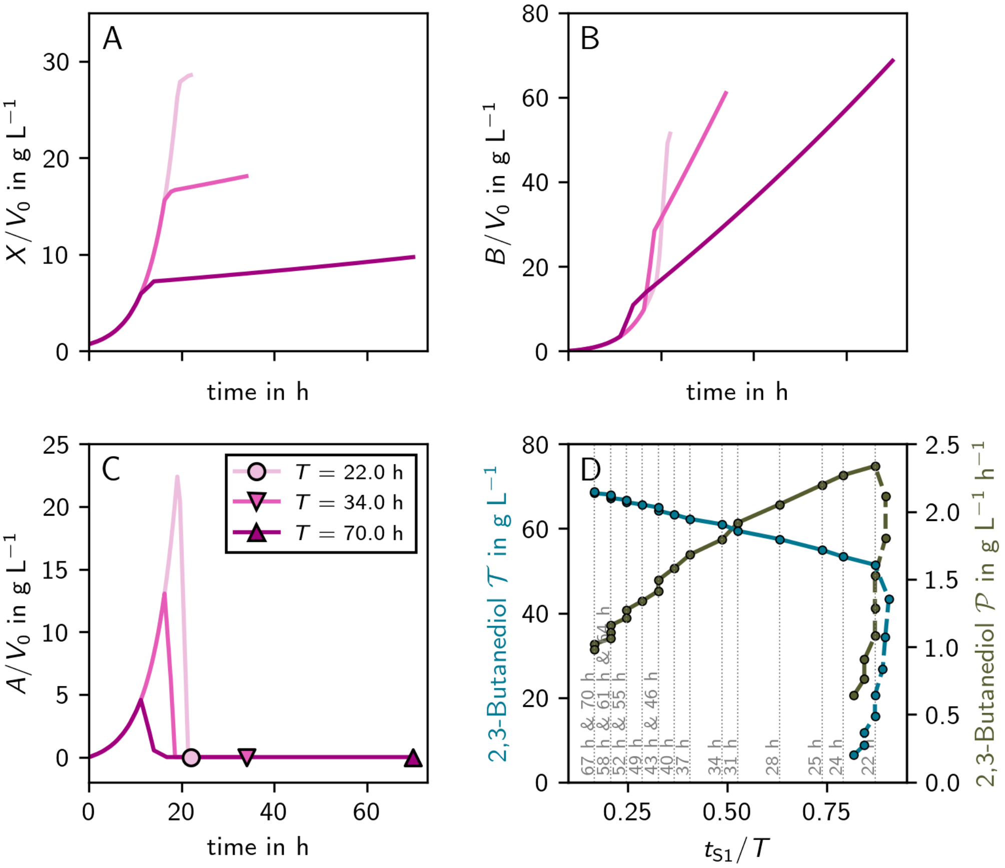


Figure S6: **Selected processes illustrating the influence of the $t_{\mathbf{S1}}/T$ ratio.** Panels A-C show the biomass, the 2,3-butanediol, and the acetoin concentrations, panel D indicates the influence of the $t_{S1}/T$ on the process metrics. Processes defined over $T$ in the legend of panel C correspond to the markers that lie on the dotted lines labeled with the same $T$ in panel D. Although the process length $T$ increases with titer $\mathcal{T}$, the relative length of the first stage $t_{S1}/T$ is reduced.

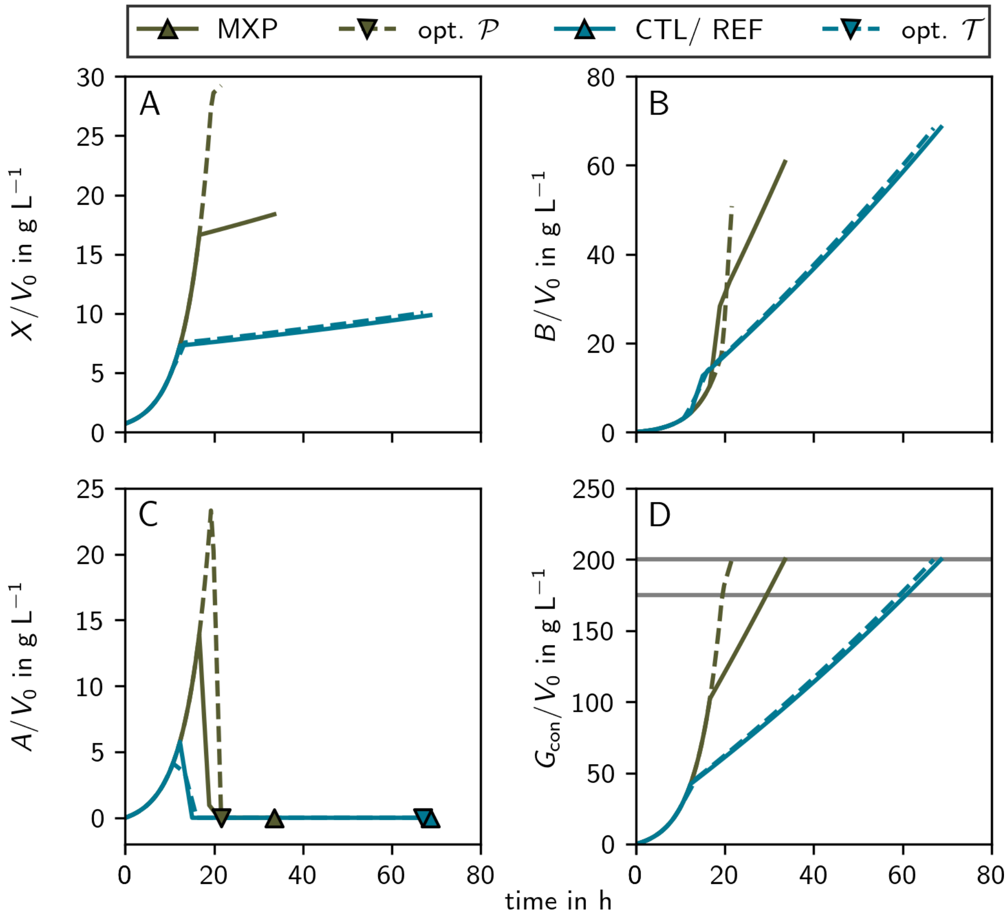


Figure S7: **Comparison between the optimal processes for productivity and titer (opt. $\mathcal{P}$ and opt. $\mathcal{T}$, respectively) shown with dashed lines and the processes as performed in the validation experiments (full lines).** While opt. $\mathcal{T}$ and CTL are very similar, there is more variation between opt. $\mathcal{P}$ and MXP. All four lines represent simulations, experimental data is shown in Figures 6 and 3. All simulations are performed until a maximum amount of $200\,\mathrm{g\,L^{-1}}$ glucose was consumed (top gray line, panel D). To compare the simulations to the validation experiments fed with a maximum of $175\,\mathrm{g\,L^{-1}}$ glucose, the time points where the simulations passed the bottom gray line (panel D) were used as the respective process ends. Both, REF and CTL, refer to the same simulation, only the end point is different, depending on the glucose fed as explained above.

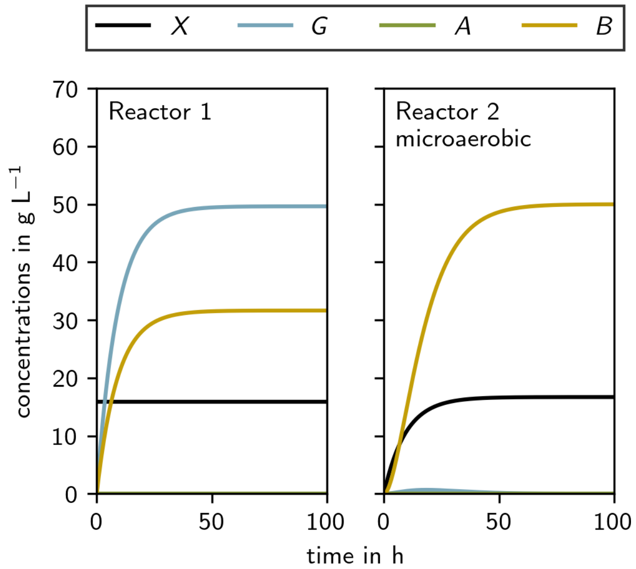


Figure S8: **Optimal productivity two-reactor continuous process simulation with a $\mathcal{T} \geq 50\,\mathrm{g\,L^{-1}}$ constraint.** Reactor A is run at $\mu = 0.11\,\mathrm{h^{-1}}$ and reactor B is run microaerobically ($\mu = 0.005\,\mathrm{h^{-1}}$). The steady state concentrations are given in Supplementary Table S2 and the process metrics are given in Supplementary Table S3.
